# Antifungal potency of terbinafine as a therapeutic agent against *Exophiala dermatitidis in vitro*

**DOI:** 10.1101/2024.05.25.595862

**Authors:** Tomofumi Nakamura, Tatsuya Yoshinouchi, Mayu Okumura, Toshiro Yokoyama, Daisuke Mori, Hirotomo Nakata, Jun-ichirou Yasunaga, Yasuhito Tanaka

**Affiliations:** Department of Laboratory Medicine, Kumamoto University Hospital, Kumamoto, Japan; Department of Hematology, Rheumatology, and Infectious Diseases, Faculty of Life Sciences, Kumamoto University, Kumamoto, Japan; Department of Medical Technology, Faculty of Health Science, Kumamoto Health Science University, Kumamoto, Japan; Department of Gastroenterology and Hepatology, Faculty of Life Sciences, Kumamoto University, Kumamoto, Japan

**Author notes:** These authors contributed equally (Tomofumi Nakamura, Tatsuya Yoshinouchi). To whom correspondence should be addressed; Tomofumi Nakamura, M.D., Ph.D., (T.N.).

**Keywords:** *Exophiala dermatitidis*, terbinafine, posaconazole, amphotericin B, biofilm

## Abstract

**Background:** *Exophiala dermatitidis* (*E. dermatitidis*), which causes skin infections or respiratory diseases, is occasionally fatal in immunocompromised patients.

**Objectives:** Here, we report the unique antifungal potency of terbinafine (TRB), which targets squalene epoxidase, against *E. dermatitidis* (SQLE^ED^) using various *in vitro* approaches.

**Methods:** Based on the human SQLE crystal structure, we created a structure model of SQLE^ED^ using SWISS-MODEL and determined the best-fitting model of TRB to the SQLE^ED^. The versatile antifungal activities, including fungicidal activity, biofilm inhibition, biofilm eradication activity, and the combination effect of TRB, posaconazole (PSC), and amphotericin B (AmB) with great antifungal potency against *E. dermatitidis* were evaluated using crystal violet and cell viability assay.

**Results:** Clinically isolated *E. dermatitidis* increased most vigorously at 30°C but decreased at 40°C. *E. dermatitidis* hyphae elongated and attached to a cell scaffold, forming a membrane-like biofilm that was distinct from the cell-free biofilm. In the binding model, TRB formed an H-bond with Y102 and was surrounded by key amino acid residues of SQLE^ED^ corresponding to TRB-resistant mutations in *Trichophyton rubrum*, showing an appropriate interaction. Among TRB, PSC, and AmB with potent antifungal activities, TRB and PSC showed more potent antibiofilm activities than AmB. In addition, TRB and PSC exhibited residual potency without incubation against *E. dermatitidis*, decreasing the growth at lower concentrations than AmB. In contrast, AmB exhibited strong time-dependent killing and eradication activities. The combination of TRB and PSC was more effective than that of TRB and AmB or PSC and AmB *in vitro*.

**Conclusions:** Although the tissue migration of TRB must be considered, these data suggest that TRB and PSC may be useful agents and a potent combination in severely immunocompromised patients with refractory and systemic *E. dermatitidis* infection.

## Introduction

It is estimated that 1.7 billion people worldwide suffer from fungal infections.^1^ Invasive fungal infections in patients undergoing organ transplantation, chemotherapy for cancer, HIV infection, or autoimmune diseases cause approximately 1.7 million deaths per year.^2^

*E. dermatitidis* is a black fungus, a member of the Herpotrichiellaceae, that can be isolated from wet living environments such as dishwashers, humidifiers, and bathtubs,^3^ and is commonly reported as a pathogen of black fungal infections isolated from the skin and subcutaneous tissue in dermatological treatment.^4^ Respiratory tract infections caused by *E. dermatitidis* are relatively rare, with reports of underlying diseases such as bronchiectasis and cystic fibrosis.^5,6^ Moreover, black fungal infections caused by *E. dermatitidis* isolated from the skin, eye, liver, central nervous system, and central venous catheters have been reported as opportunistic fatal infections in immunocompromised patients.^7^ *E. dermatitidis* has a slower growth rate than the other fungi. Small black colonies of *E. dermatitidis* observed on Sabouraud dextrose agar (SDA) after 3 days were barely detectable. Matrix-assisted laser desorption ionisation-time of flight mass spectrometry (MALDI-TOF MS) is a powerful tool for the early identification of pathogenic fungal species, including late-growing fungi such as *E. dermatitidis*, in addition to morphological and genetic analysis of the specimen.^8^ We previously reported *E. dermatitidis* pneumonia in immunocompromised patients with anorexia nervosa.^9^ In clinical practice, *E. dermatitidis* is treated with azoles such as voriconazole and itraconazole for several months.^10,11^ In this study, we analysed characteristics of *E. dermatitidis* and evaluated the antifungal, antibiofilm, killing (fungicidal) activity, and combinations of several clinically used oral and intravenous antifungal agents such as terbinafine (TRB)^13^, azoles (fluconazole [FLC], miconazole [MCZ], voriconazole [VRC], itraconazole [ITC], PSC, isavuconazole [ISC]),^12^ micafungin (MCF), caspofungin (CAS), and amphotericin B (AmB)^14^ against *E. dermatitidis*. TRB, an allylamine medicine, showed a potent and unique anti-fungal effect against *E. dermatitidis*, which could lead to better treatment options that may prevent fatal *E. dermatitidis* infection in immunocompromised patients.

## Material and Methods

### Fungus and Cells

*E. dermatitidis* 1 and 2 isolated from the patients with pneumonia were identified by ESI-MS and ITS gene analysis^15^, and *E. dermatitidis* 3 (NBRC6421, ATCC28869) was purchased from Biological Resource Center, NITE (NBRC, Japan) (Table S2). The fungus was incubated in Sabouraud buffer (5 g meat peptone, 5 g casein peptone, and 20 g glucose in 1L dH_2_O) and Sabouraud dextrose agar (SDA) plate (5 g meat peptone, 5 g casein peptone, 40 g glucose, and 1.5% agar in 1L H_2_O) supplemental with chloramphenicol (Cam) and kanamycin (K), or 0.25µm filtered RPMI (Nissui, Japan) without NaHCO_3_ at pH 6.8, 0.165M 3-morpholinopropane-1-sulfonic acid, MOPS (MOPS-RPMI) supplemental with Cam and K. A549 cells isolated from a male patient with lung cancer were purchased from the Japanese Collection of Research Bioresources (JCRB) Cell Bank, Japan, and cultured in DMEM medium (FUJIFILM Wako Pure Chemical Corporation, Japan) supplemented with 10% fetal bovine serum (FBS; Gibco, Thermo Fisher Scientific, USA), penicillin (P), and K.

### DNA and RNA extraction, and identification of SQLE sequences

A collection of *E. dermatitidis* incubated on SDA with a small medicine spoon was completely frozen in liquid nitrogen. After grinding with a masher tube, total RNA and DNA were extracted using TRIzole (Invitrogen, Thermo Fisher Scientific). RNA was converted into cDNA using ReverTra Ace® (TOYOBO, Japan). These primers (ITS4R: TCC TCC GCT TAT TGA TAT GC NS7F: GAG GCA ATA ACA GGT CTG TGA TGC) provided the best combination for identification with the strain of *E. dermatitidis* using the ITS sequence. PCR amplifications were performed in an Eppendorf thermocycler (Eppendorf® Mastercycler) in a final volume of 40 μL with 10–50 ng of the DNA as a template using KOD one polymerase (TOYOBO). The SQLE amino acids sequence of *E. dermatitidis* was identified from the cDNA using these primers (ED.SE.SF: ATG CCT CTC ATA CTC GAT TCG TCG TC, ED.SE.ER: TCA AAT CCT CAG TTC GGC AAA TAT ATA CG, ED.SE.SQF: TCT GAT TCT GGG TGT GGA GTC C, ED.SE.SQR: TCA GGT ACG TCG ACC AGG ACA CG)

### SQLE 3D structure and docking Simulation

The SQLE^ED^ sequence was identified from the genomic DNA and RNA of *E. dermatitidis*. The 3D structure model of SQLE^ED^ was produced as a template of the crystal structure of SQLE^Hum^ (PDB accession number, 6C6N) using the SWISS model (https://swissmodel.expasy.org/). A nicotinamide adenine dinucleotide (NAD) was docked to the SQLE^ED^ in the same manner as the crystal structure of SQLE^Hum^ using SeeSAR v13.1 (BioSolveIT GmbH, Sankt Augustin, Germany).^48^ Next, we determined the binding pocket of TRB using the 6C6N and clinically isolated TRB-resistant amino acid mutations of *T. rubrum* and performed the docking simulation of TRB to the SQLE^ED^. Molecular graphics and analyses were performed using UCSF Chimera (https://www.rbvi.ucsf.edu/chimera).

### Minimum Inhibitory Concentration

According to CLSI M38 3rd edition, *E. dermatitidis* was inoculated at 1.0 × 10^5^ CFU/ml and incubated at 35℃ for 48 or 72 hours in the flat-bottomed, 96-well plate with MOPS-RPMI. MIC (mg/L) represents the lowest concentration of antifungal agents that inhibited the visible growth of *E. dermatitidis*. MIC_50_ and MIC_90_ (mg/L) are determined by measuring fungal growth at OD_530_ nm or proliferation of living cell at OD_440_ nm after staining 2-(4-Iodophenyl)-3-(4-nitrophenyl)-5-(2,4-disulfophenyl)-2H-tetrazolium, monosodium salt, WST-1 (Dojindo, Japan) using an absorbance spectrometer (FLUOstar Omega, BMG Labtech, Germany).

### Drug combination

Drug combination assay referred to the previous report^25^. Briefly, combinations between TRB and azoles, AmB and azoles, TRB and AmB, or TRB and CAS at tested concentration were prepared in a polystyrene, flat-bottomed, 96-well plate with MOPS-RPMI medium. The conidium of *E. dermatitidis* was seeded at 1.0 × 10^5^ CFU/mL and incubated at 35℃ for 48 or 72 hours. MIC (mg/L) was determined by the visible growth of *E. dermatitidis*. MIC_50_ and MIC_90_ (mg/L) were calculated by measuring fungal growth at OD_530_ nm or proliferation of living cells at OD_440_ nm after WST-1 staining. The FIC index (FICI) of tested combinations was identified according to previous reports^25,49^. Synergy effect decided FICI < 2.0, no interaction; 2.0 < FICI < 4.0, and antagonistic effect; FICI > 4.0. from each MIC, MIC_50_ (WST-1), and MIC_90_ (WST-1) data.

### Time-kill assay

The time-kill assay of *E. dermatitidis* (1.0 × 10^6^ CFU/ml) was performed in a microtube tube with 100 μL of MOPS-RPMI medium. TRB, AmB, and PCZ were tested at different concentrations ranging from 0.13 to 32 mg/L. The samples were shaken at 200 rpm, and incubated at 35°C for 0, 3, 6, and 12 hours, respectively. The samples with or without drugs at various concentrations were washed twice with 1000 μL phosphate-buffered saline (PBS) at centrifugation of 2500 rpm for 2 minutes. The *E. dermatitidis* samples were duplicated (final concentration, 2.5 × 10^5^ CFU/ml) and inoculated into a 96-well plate with new 200 μL of MOPS-RPMI medium without drugs at 35°C for 48 hours. The viable *E. dermatitidis* was evaluated by the WST-1 staining assay.

### Observation of *E. dermatitis* morphology with or without A549 cells

A549 cells were used to observe the adhesion ability of *E. dermatitis* by microscopy and Scanning Electron Microscope (SEM), JSM-IT300 InTouchScope™.^9,50^ A549 cells at 1.5 × 10^5^/ml were inoculated into a 96-well plate or a coverslip, 18 × 18 mm, set on a chamber slide Ⅱ (IWAKI, Japan), filled with DMEM containing 10% FBS, P, and K. After O/N incubation in 5% CO_2_ at 37℃, the final concentration of *E. dermatitis* was added at 1.0 × 10^6^/mL to these plates, which were changed to DMEM containing 1% FBS, P, and K, and incubated in 5% CO_2_ at 35℃. The tested drugs at 0.25 mg/L were added to the plate after 6 hours. These samples were further incubated in 5% CO_2_ at 35℃ for 24 or 48 hours. The A549 cells and *E. dermatitis* in the 96-well plate were observed under a microscope after May-Giemsa staining. The samples on the coverslip in the chamber slide Ⅱ were fixed with 2% glutaraldehyde in 0.1 M phosphate buffer (pH 7.4) for 24 or 48 hours and dehydrated through 50, 75, 90, 95, and 100% ethanol sequentially. The 100% ethanol was replaced with t-butyl alcohol to cover the samples and stored at −20℃. The frozen sample was lyophilized under a vacuum and was coated with platinum. SEM observation was performed.

### Biofilm Inhibition and Eradication

Biofilm inhibition was determined using the crystal violet (CV) staining assay^23^. The conidium of *E. dermatitidis* was seeded at 1.0 × 10^5^ CFU/ml in a flat-bottomed, 96-well plate with the tested drugs and incubated at 35℃ for 48 hours. The antifungal activity of the tested drugs was also determined by measuring OD_530_. The samples were washed twice with 200 μL PBS, and stained with 100 μL of a 0.1% CV solution for 20 minutes at room temperature. The samples were washed again with 200 μL PBS and dried overnight. The 100 μL of 30% acetic acid was incubated for 30 minutes to extract CV staining from the biofilm. The 80 μL of the solution was transferred to a fresh 96-well plate and the samples were measured at OD_620_ nm. Biofilm inhibition by the drugs was expressed as a relative ratio compared to positive controls (no drugs). A modified biofilm eradication assay was performed according to the previous report^51^. Briefly, the conidium of *E. dermatitidis* was seeded at 5.0 × 10^5^ cells/ml in a 96-well plate without the tested drugs in 100 μL of MOPS-RPMI and incubated at 35℃ for 24 hours. Then, the tested drugs were added to the plate at each concentration and further incubated at 35℃ for 24 hours. The eradication ability of the tested drugs was determined by the CV staining assay described above.

### Illustration

Illustrations accompanying the experimental procedure and explanations were created using BioRender: Scientific Image and Illustration Software.

### Drugs

TRB, MCZ, VRC, ITC, and PSC were purchased from the Tokyo Chemical Industry Japan, FLC and AmB from Wako Japan, ISC and CAS from Selleck Chemicals USA, and MCFG from Cayman Chemical USA. These tested drugs (5–20 mM) in DMSO or appropriate solutions were stored at −80℃. Before the assays, these drugs were prepared at appropriate concentrations in MOPS-RPMI buffer.

## Results

### Profile of clinically isolated *E. dermatitidis*

We observed giant black colonies of *E. dermatitidis* 1 isolated from a clinical patient^9^ under aerobic incubation at room temperature on SDA and performed morphological observations using an optical microscope and scanning electron microscope (SEM) (Fig. 1). The centre of the giant colonies formed a black gelatinous mass and the surface and edges changed to olive brown (Fig. 1A). Microscopy revealed many yeast-like oval conidia in the centre (Fig. 1B) and septate hyphae and hyphae in the outer part of the black colony (Fig. 1B). The internal transcribed spacer 1 (ITS1) and D1/D2 regions of the RNA were compared using the Basic Local Alignment Search Tool (BLAST). The identification rate of the D1/D2 region was dominated by *Exophiala* species, and that of *E. dermatitidis* was 96.9%, and the ITS region was 100% identical to *E. dermatitidis* (Fig. S1A). Based on genetic analysis of the ITS region, we identified the giant black fungus as E. *dermatitidis* genotype A (Fig. S1B), which is most commonly reported in Japan.^15^

**Figure 1.**
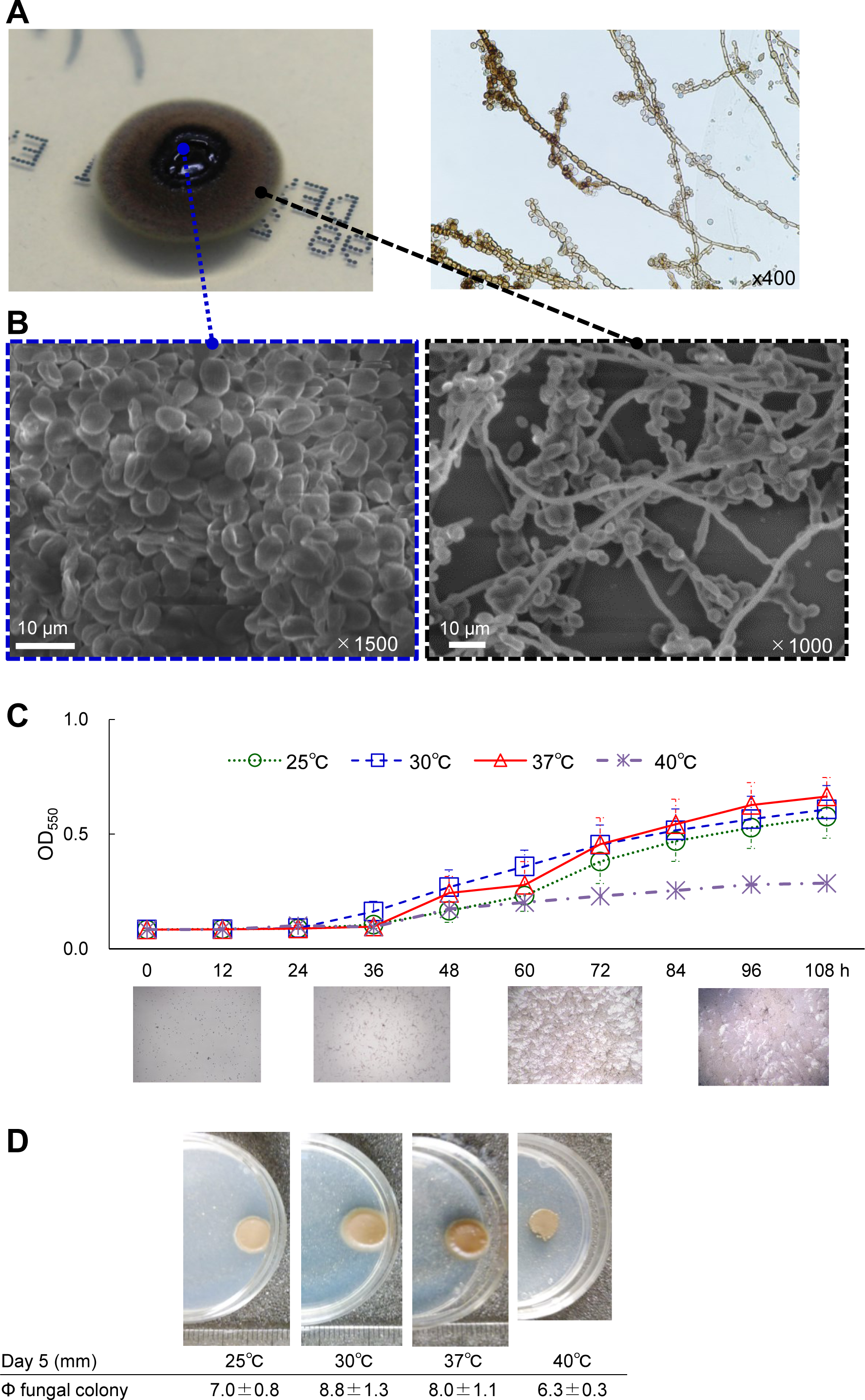
Characteristics of *E. dermatitidis*. **A**) *E. dermatitidis* colony incubated on an SDA plate for 14 days was black gelatinous in the centre, changing to brown at the edges. The lactophenol cotton blue-stained specimen image was observed by the slide culture method. **B**) Morphological image of the central and marginal areas of *E. dermatitidis* colony by SEM. The scale bar and magnification at each image are shown **C**) Growth rate and microscopic images of *E. dermatitidis* in MOPS-RPMI culture medium until 108 hours or **D**) colonies on the SDA plate for 120 hours at 25, 30, 37, and 40℃, respectively.

Next, we analysed the growth of *E. dermatitidis* at 25, 30, 37, and 40℃. The growth increased more efficiently at 30 and 37℃ than at 25 and 40℃ in MOPS-RPMI (Fig. 1C) and Sabouraud medium (Fig. S1C). In addition, the diameter of the black colonies of *E. dermatitidis* on SDA incubated for a week was 8.8, 8.0, 7.0, and 6.3 mm at 30, 37, 25, and 40℃, respectively, in line with the growth results at these temperatures (Fig. 1D). These results suggest that the growth of *E. dermatitidis* was affected by incubation temperature; in particular, incubation at 40℃ prevented the growth. The benefit of thermotherapy in cutaneous or subcutaneous infection has been previously reported.^16^

### Structural modelling of *E. dermatitidis* squalene epoxidase

Target proteins of various antifungal drugs in clinical use are shown in Fig. 2. TRB target squalene epoxidase (SQLE), also known as squalene monooxygenase, which catalyses the conversion of squalene to (S)-2,3-epoxy squalene, is a key enzyme in ergosterol biosynthesis in fungal membranes.^17,18^ Azoles can also exert anti-fungal activity by inhibiting a different target enzyme, lanosterol 14α-demethylase in the same pathway as shown in Fig. 2A.^19^

**Figure 2.**
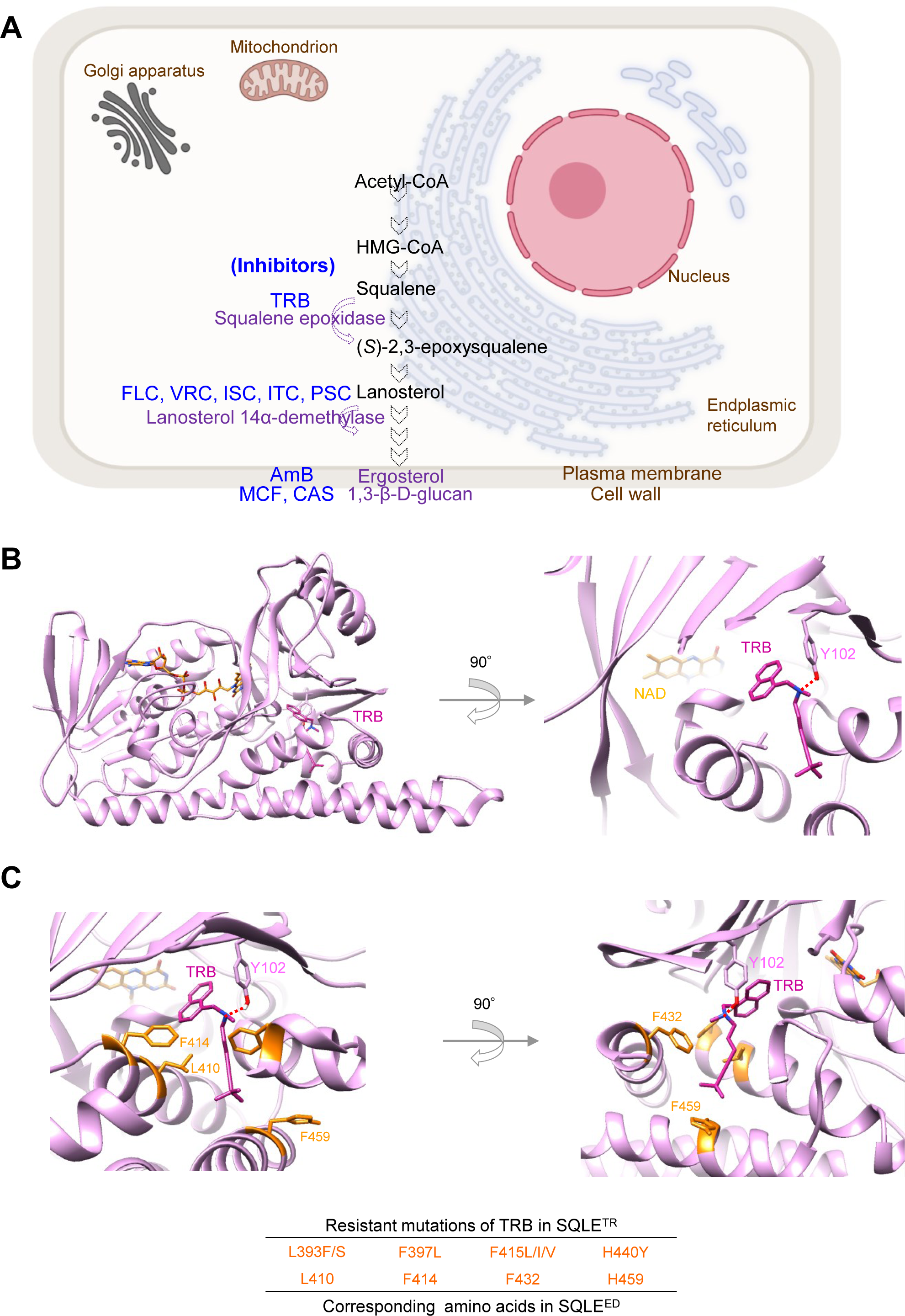
Targets of antifungal drugs and SQLE structures. **A**) Schematic illustration of antifungal target proteins in fungal cells. **B**) The best binding model of TRB to the SQLE structure of *E. dermatitidis* with nicotinamide adenine dinucleotide (NAD). TRB interacts with Y102 through an H-bond in SQLE^ED^. **C**) The location of putative TRB-resistant amino acid residues (L410, F414, F432, and H459) in the SQLE^ED^ structure corresponding to clinically TRB-resistant mutations (L393F/S, F397L, F415L/I/V, and H440Y) in the SQLE^TR^.

To investigate the structural antifungal efficacy of TRB against *E. dermatitidis*, we identified the SQLE^ED^ derived from three *E. dermatitidis* (Table S1) with the same amino acid (AA) sequences. The AA sequences of SQLEs from humans, *E. dermatitidis*, and *Trichophyton rubrum* were compared using Clustal omega (multiple sequence alignment tool), as shown in Table S1. Moreover, the structural modelling of SQLE^ED^ was produced by SWISS-MODEL based on the crystallography of human SQLE (SQLE^Hum^) (PDB:6C6N) as previously reported^18^ and the full-length SQLE^Hum^ model shown in Fig. 2B and S2A. Next, we performed a docking simulation of TRB to SQLE^ED^ (Fig. 2B) based on the crystal structure of Cmpd-4 bound to SQLE^Hum^ (PDB:6C6N) using SeeSAR (see Methods). The top 20 structures of the TRB bound to SQLE^ED^ are shown in Fig. S2C. The TRB-binding structures of SQLE^Hum^ and SQLE^ED^ formed hydrogen bonds with the side chains of Tyr 122 and 102, respectively (Fig. S2). Important AA residues (L393, F397, F415, and H440) of SQLE (SQLE^TR^) that bind to TRB have been reported in the resistance profiles of clinical isolates of *T. rubrum* (Fig. S2D).^31^ TRB is surrounded by L410, F414, F432, and H459 of SQLE^ED^, which correspond to L393, F397, F415, and H440 of SQLE^TR^ in the biding respectively (Fig. 2C). These results indicate that TRB can effectively interact with SQLE^ED^, resulting in a potent antifungal effect against *E. dermatitidis*.

### Antifungal activity of clinically used drugs against *E. dermatitidis*

Various orally and intravenously administered antifungal drugs are available for clinical use. In this study, we evaluated the antifungal effects of ten drugs (FLC, MCZ, VRC, ITC, PCZ, ISC, TRB, AmB, CAS, and MCF) (Fig. S3) against clinically isolated *E. dermatitidis* 1 (Table 1) based on the CLSI-modified M38Ed3. Additionally, the susceptibilities of the other two *E. dermatitidis* strains (E.D.2, clinically isolated, and E.D.3, ATCC28869) to TRB, PSC, and AmB were evaluated (Table S2). The minimum inhibitory concentration (MIC), 50% inhibition of growth (MIC_50_), and 90% inhibition of growth (MIC_90_) were determined using OD_530,_ or OD_440_ after WST-1 reagent staining to evaluate the viability of *E. dermatitidis* (WST-1 assay).^20^ The MICs of FLC, MCF, and CAS against *E. dermatitidis* exceeded 16 mg/L as previously described.^21,22^ PSC, VRC, AmB, TRB, and ITC showed potent inhibitory ability from 0.031 to 0.25 mg/L at MIC_50_ and MIC_90_ as assessed using OD_530_ and WST-1. MCZ at 0.25 to 0.5 mg/L inhibited the growth similarly to ISC. The MIC of these drugs was very close to MIC_90_ obtained from WST-1. *E. dermatitidis* exhibited different growth rates at different temperatures (Fig. 1C and S1C). To examine the effect of the incubation temperature on antifungal activity, we determined MIC_50_ and MIC_90_ of drugs by measuring *E. dermatitidis* growth at OD_530._ (Table S3). The findings indicated that incubation temperature did not significantly affect the antifungal activity of these drugs.

### Inhibition and eradication of *E. dermatitidis*-induced biofilm

Bacterial and fungal biofilms form in infected organs, particularly on medical devices such as intravascular catheters, artificial heart valves, and artificial joints, causing intractable chronic infections that are resistant to antibiotics and antifungals.^22^ It has been reported that *E. dermatitidis* can produce biofilm formation.^23^

*E. dermatitidis* was incubated with or without A549 cells in a glass-bottomed slide chamber plate. *E. dermatitidis* without A549 cells floated in the buffer, whereas *E. dermatitidis* with A549 cells firmly attached to the bottom of the slide (Fig. S3). Next, we investigated the morphology of *E. dermatitidis* with or without A549 cells by optical microscopy (Fig. 3A) and SEM. (Fig. 3). The coadunate filamentous biofilm-like morphology of *E. dermatitidis* was sparsely observed without A549 cells as observed by SEM (Fig. S3). On the other hand, *E. dermatitidis* hyphae with the cells extended cohesively below and above the cells, and the oval conidia were diffusely attached to the cells for 24 hours of incubation (Fig. 3B). Moreover, membrane biofilm-like morphology including the cells appeared where *E. dermatitidis* was highly enriched for 48 hours of incubation (Fig. 3C and S3). Treatment with TRB, PSC, and AmB at 0.25 mg/L against *E. dermatitidis* inhibited hypha growth and showed pseudohyphae (Fig. 3D). There were no clear differences in *E. dermatitidis* morphology between these drug treatments.

**Figure 3.**
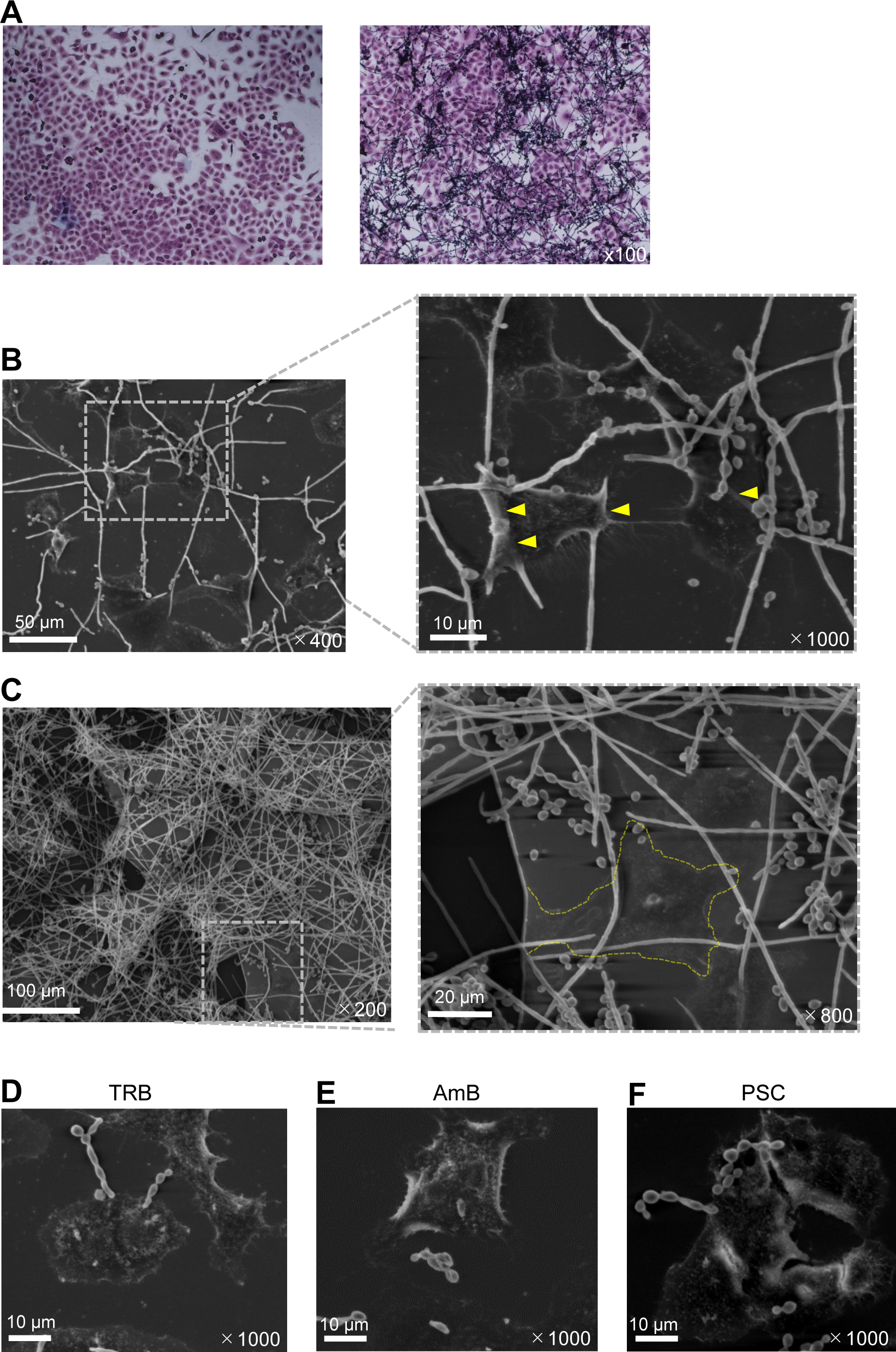
Morphology of *E. dermatitidis* with A549 cells. **A**) May-Giemsa staining on A549 cells. *E. dermatitidis* was observed by optical microscopy incubated at 35℃ for 24 hours after addition to the A549 cells. **B**) *E. dermatitidis* hyphae were elongated and attached to the 549 cells incubated at 35℃ for 24 hours by SEM. **C**) Gray-membrane biofilm containing A549 cells circled at the dotted yellow line formed on enriched *E. dermatitidis* after incubation at 35℃ for 48 hours. **D**) *E. dermatitidis* incubated on the A549 cells at 35℃ for 24 hours in the presence of TRB, AmB, and PSC at 0.25 mg/L. The scale bar and magnification at each image are shown in white.

Next, we examined the inhibitory and eradication abilities of TRB, PSC, and AmB against *E. dermatitidis*-induced biofilms using a CV assay (Fig. 4 and S4). Interestingly, TRB and PSC showed more potent inhibitory activity against the biofilm formation by *E. dermatitidis* (Fig. 4B), and the antibiofilm activity of these drugs was similar to their anti-fungal activity (Fig.S4). In contrast, AmB sufficiently and TRB slightly at high concentration eradicated the *E. dermatitidis*-induced biofilm compared to PSC (Fig. 4C). These results indicate that TRB and PSC can inhibit biofilm formation at lower concentrations than AmB. However, a higher concentration of AmB can moderately eradicate the biofilm formed, suggesting that TRB and PSC may be useful in the early treatment of acute *E. dermatitidis* infection and that AmB may be effective in the late or prolonged treatment of chronic biofilm-forming infections.

**Figure 4.**
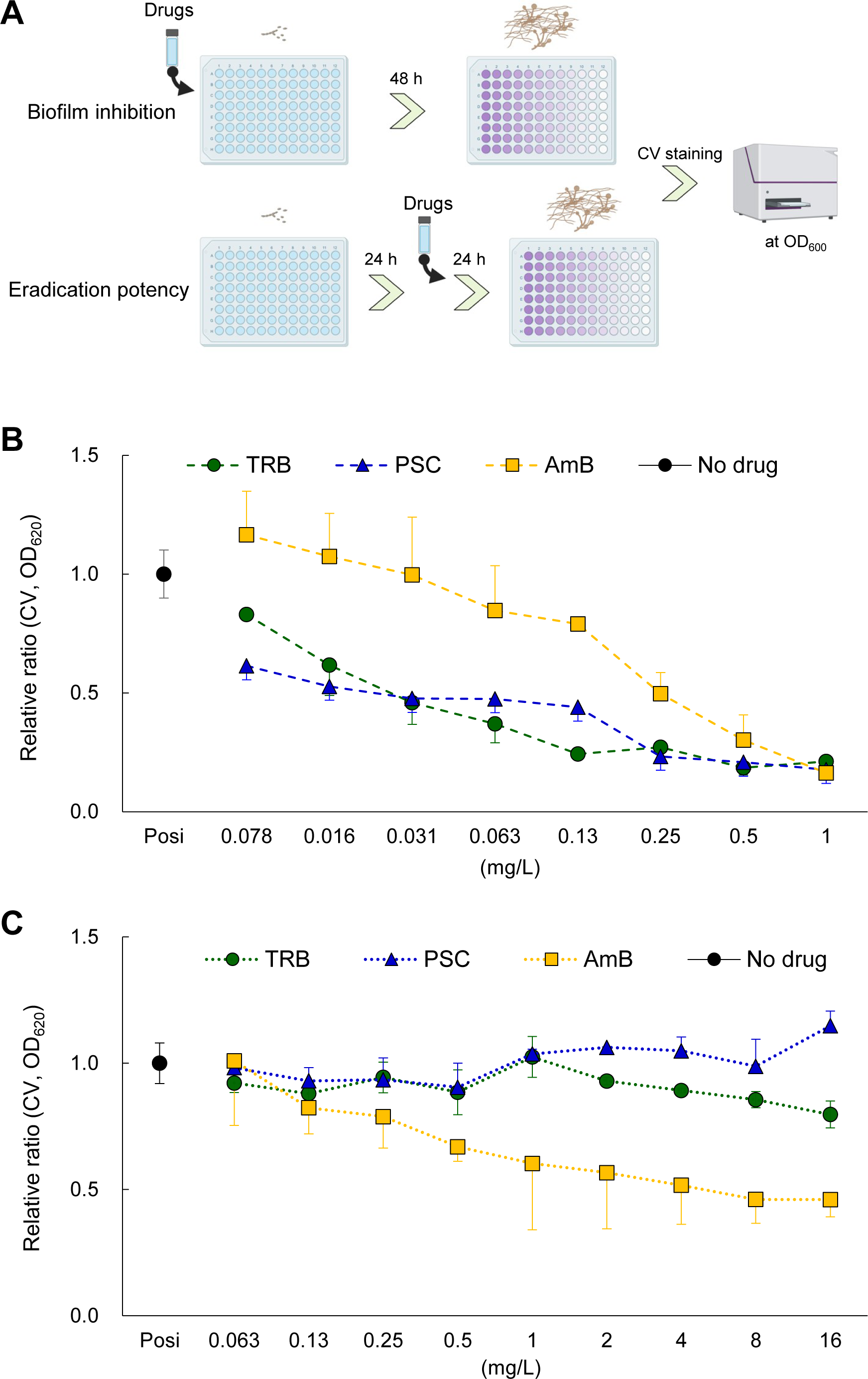
Inhibition and eradication of biofilm activity of TRB, PSC, and AmB. **A**) Schematic illustration shows assay procedures to confirm inhibition and eradication of TRB, PSC, and AmB against *E. dermatitidis* biofilm. **B**) Biofilm inhibition and **C**) eradication potency of TRB (green circle), PSC (blue triangle), and AmB (yellow square) are shown as a relative ratio (positive control is no drug) from 0.078 to 1 mg/L and from 0.063 to 16 mg/L, respectively. All assays were performed independently in triplicate. Means (± S.D.) of all data are presented.

### Fungicidal activity and combination effects of antifungal drugs against E. *dermatitidis*

The ability to kill invasive fungi is important in immunocompromised conditions.^24^ We examined a time course of the killing ability of these drugs using WST-1 reagent to examine living *E. dermatitidis*. Notably, TRB, PSC, and AmB could be inhibited or killed by temporal attachment (no incubation) to *E. dermatitidis* conidia (see Materials and Methods). (Fig. 5A and B). TRB and PSC at lower concentrations (from 0.5 mg/L) decreased the growth of *E. dermatitidis*, indicating that TRB and PSC rapidly interact with and maintain the target proteins. In addition, during the observation of the killing ability of the drugs from 0 to 12 hours, TRB did not show any killing ability even at 32 mg/L. The viability of *E. dermatitidis* incubated for 12 h at all TRB-tested concentrations increased compared to that without incubation, suggesting that *E. dermatitidis* can withstand and show a reactive response to TRB pressure (Fig. 5C). PSC and AmB killed *E. dermatitidis* in dose- and time-dependent manner (Fig. 5D and E).

**Figure 5.**
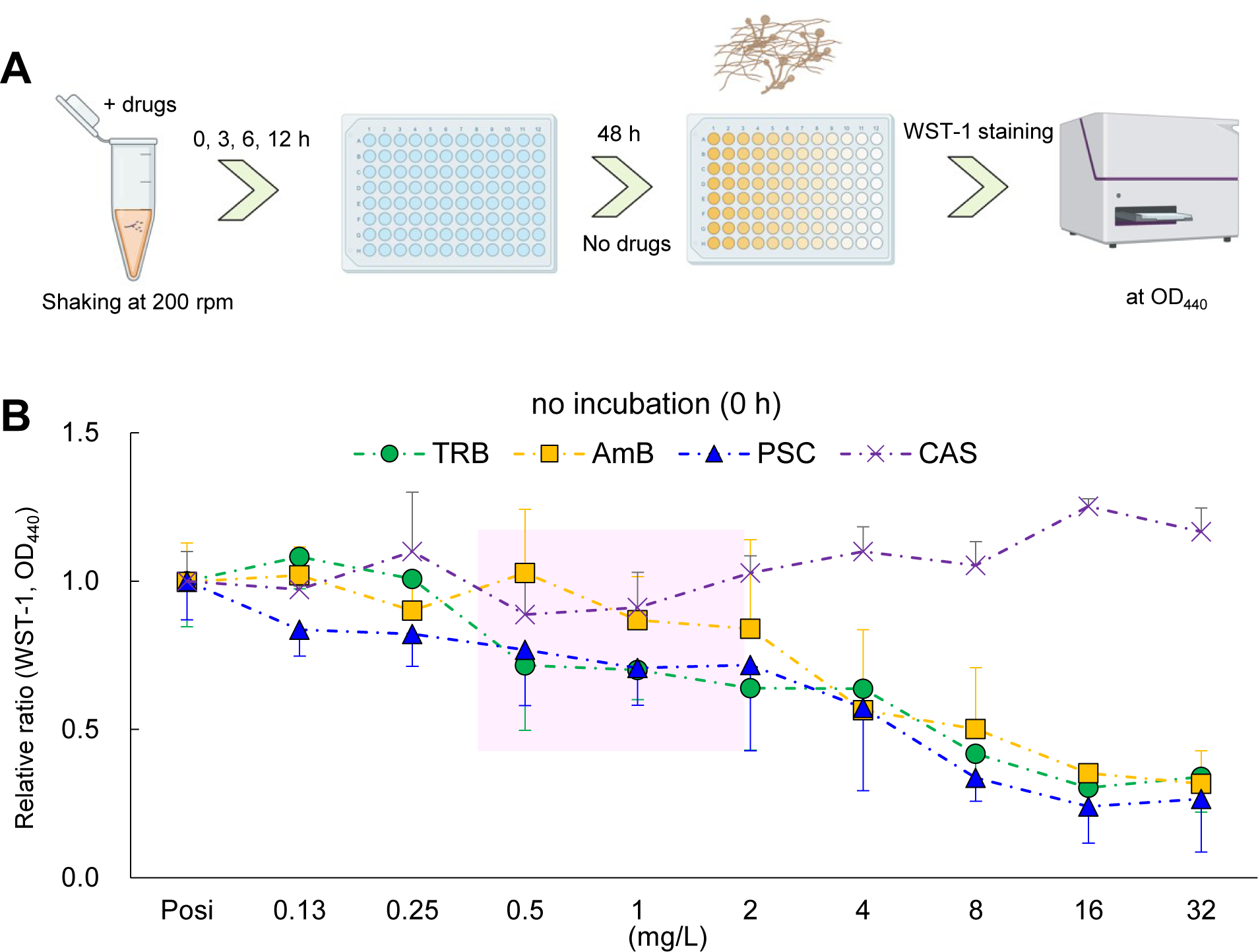
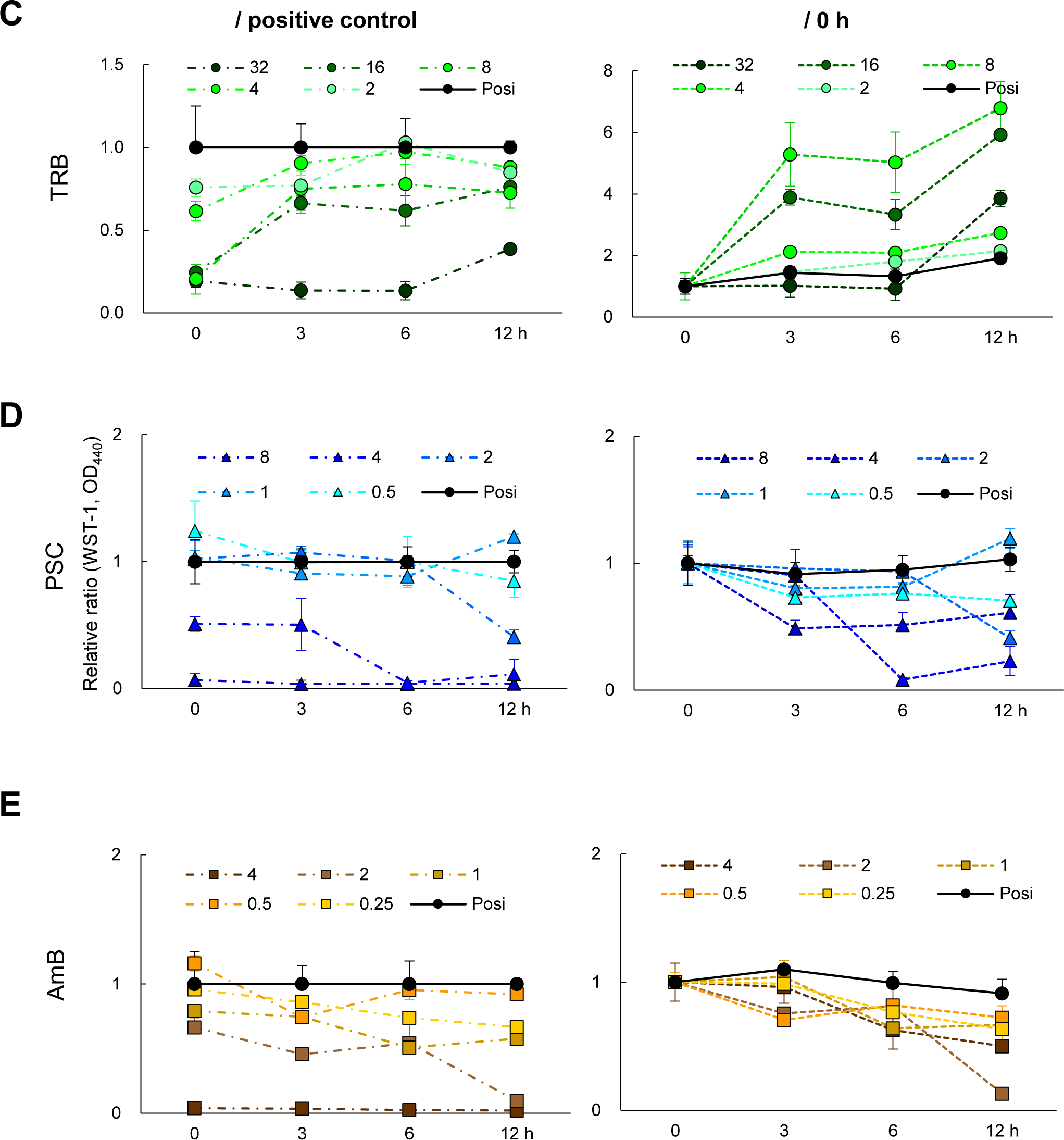
Residual and killing activity of TRB, PSC, and AmB. **A**) Schematic illustration of assay procedures to confirm the residual and killing potency of TRB, PSC, and AmB against *E. dermatitidis*. **B**) Residual potency of TRB (green circle), PSC (blue triangle), AmB (yellow square), and CAS (purple × mark) treated with no incubation time (0 h) is shown as a relative ratio (positive control is no drug) from 0.125 to 32 mg/L. Assays were performed independently in triplicate. Means (± S.D.) of all data are presented. The killing potency of **C**) TRB, **D**) PSC, and **E**) AmB were evaluated and shown as relative ratios at each appropriate concentration and incubation time as assessed by cell viability using WST-1 staining. Assays were performed independently in triplicate. Means (± S.D.) of representative data are shown.

*In vitro* and *in vivo* combination therapies are effective in treating refractory and chronic infections.^25^ TRB and other antifungal combinations have been reported.^26^ We investigated the combinations of antifungal drugs such as TRB and azoles (PSC, VRC, and ITC), TRB and AmB, and TRB and CAS against *E. dermatitidis* (Fig. 6 and S5). A previous study^21^ showed a synergistic effect against *E. dermatitidis* when CAS was combined with azoles (VRC and ITC) *in vitro*. The fractional inhibitory concentration index (FICIs) was calculated from the MIC, MIC_50_, and MIC_90_ values in the WST-1 assay. The values for CAS and PCZ were < 0.5, indicating a synergistic effect (Fig. S5). The value for the combination of TRB and AmB was between 1.0 and 2.0, and that for PCZ and AmB between 1.0 and 2.2, resulting in no interaction effects (Fig. 6). However, when a combination of TRB with other azoles was examined, the FICI of TRB and PSC showed better efficacy between 0.31 and 0.75, TRB and ISC between 0.63 and 1.0, TRB and ITC at 0.5, and TRB and VRC between 0.63 and 1.0, indicating synergistic or no interaction effects (Fig.6 and Fig. S5). These results suggest that the combination of TRB with azoles that inhibit different target proteins in the same pathway,^26^ particularly ITC and PCZ, was more favourable than AmB against *E. dermatitidis*.

**Figure 6.**
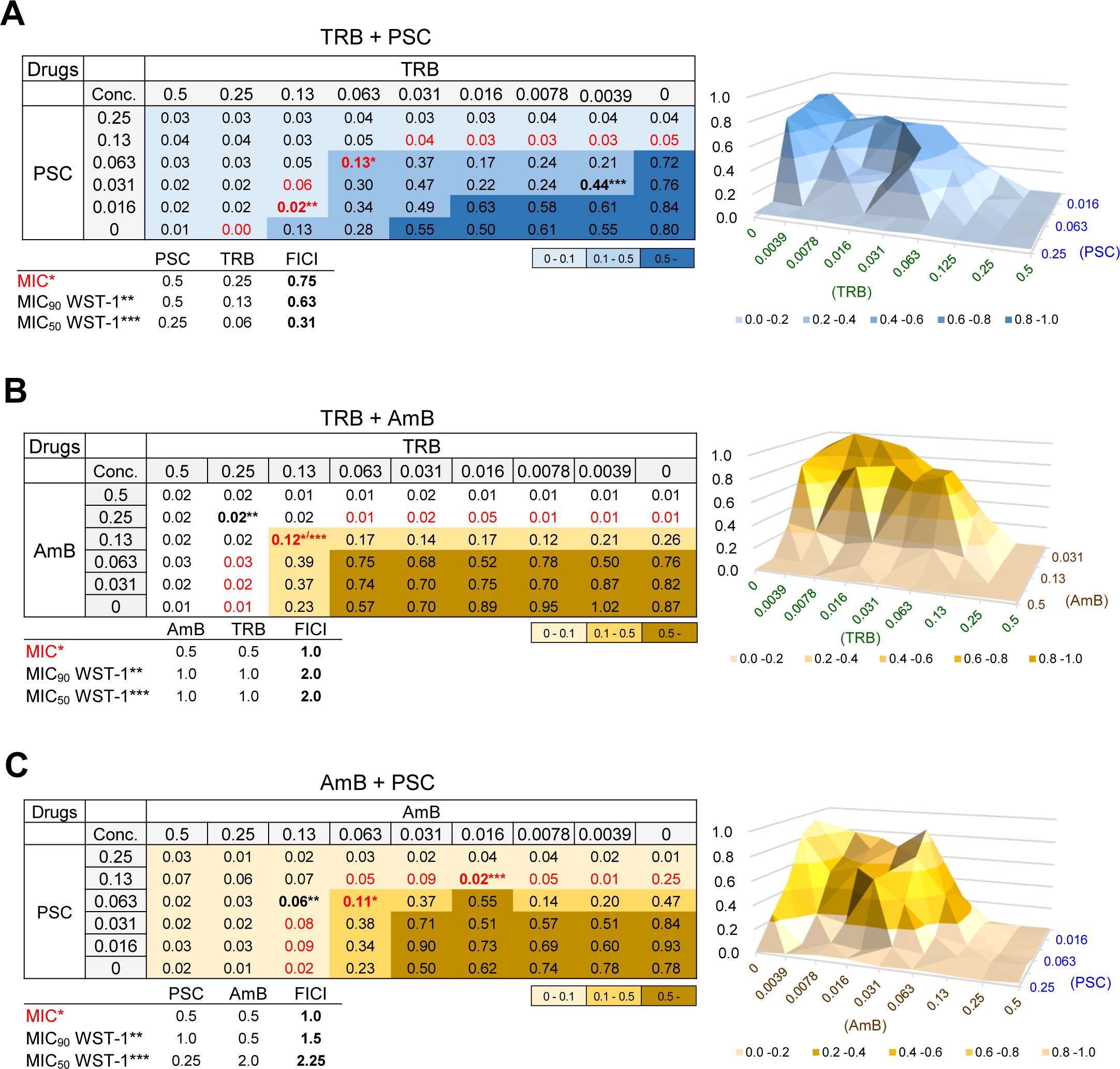
Combination effect of TRB, PSC, and AmB on *E. dermatitidis*. A combination of **A**) TRB and PSC, **B**) TRB and AmB, and **C**) AmB and PSC tables and 3D surface plots consist of relative ratios at each appropriate concentration as assessed by *E. dermatitidis* viability using WST-1 staining. FIC indices (FICI) were evaluated as synergy; FICI < 0.5, no interaction; 0.5 < FICI < 4, or antagonism; FICI > 4, which calculated from MIC (*in red) data or MIC_50_ (***) and MIC_90_ (**) of WST-1 staining results. All assays were performed independently in triplicate and representative data are shown.

Taken together, TRB may be a therapeutic agent with novel antifungal, anti-biofilm, and residual activities against *E. dermatitidis* in combination with azoles *in vitro* (Table 2).

## Discussion

Systemic and invasive *E. dermatitidis* infections such as pneumonia and sepsis require a relatively long treatment period,^27^ and have been reported to result in death.^8^ Therefore, potent antifungal activity and tissue migration of antifungals to the site of infection are essential for antifungal treatment when treating immunocompromised patients.

In this study, TRB showed potent antifungal activity against *E. dermatitidis* similar to that of PSC in various *in vitro* assays. In addition, TRB showed a stronger anti-biofilm effect (Fig. 4A) than AmB, similar to PSC against fungus biofilm formation that is resistant even to disinfectants.^28^ The residual potency of TRB for the treatment of *E. dermatitidis* does not require a higher effective drug concentration for a longer time (Fig. 4C), indicating that TRB exerts a dose-but not time-dependent effect on *E. dermatitidis* in terms of pharmacokinetic-pharmacodynamic parameters.^29^

TRB that targets SQLE is used in clinical practice as an oral medication, cream, and solution for the treatment of dermatomycoses such as *trichophytosis*.^30,31^ The important AA residues (L393, F397, F415, and H440) of SQLE that bind to TRB have been reported in the resistance profiles of clinical isolates of *T. rubrum*.^32^

We identified the AA residues of SQLE^ED^ from the genome and confirmed that the AA residues at L410, F414, F432, and H459 of SQLE^ED^ corresponded to those at L393, F397, F415, and H440 of SQLE^TR^, respectively, indicating that there were no TRB resistance mutations against *E. dermatitidis* (Table S1 and S2). In the binding model, TRB formed an H-bond with Y102 and was surrounded by TRB resistance-related residues at L410, F414, F432, and H459, showing a better interaction (Fig.2 and Fig. S2). These data suggested that TRB can strongly bind to SQLE^ED^, eliciting a potent antifungal effect.

TRB shows sufficient serum concentrations^33,34^ and drug transfer to the skin and nails,^35^ resulting in efficacy against dermatomycosis, but insufficient transfer to lung tissue^36,37^ in animal models. Nevertheless, Oral administration of TRB has been reported to be effective in a few patients with chronic *Aspergillus* lung infections.^38,39^ Clinical trials with larger numbers of patients are needed to evaluate the efficacy of TRB in the treatment of fungal pneumonia.

We believe that TRB prophylaxis or treatment of sepsis and infected sites with high blood flow may be effective, because TRB is a drug with sufficiently elevated blood levels, anti-biofilm, and potent residual effects. In contrast, direct administration of TRB as an external drug or by inhalation may be preferable in areas with poor blood flow and transitivity, such as intrabronchial fungal infections.

In this study, PSC showed a potent antifungal activity profile that was different from that of TRB against *E. dermatitidis*. According to a report on tissue concentrations in biopsy specimens obtained at autopsy from seven patients receiving PSC prophylaxis, lung concentrations were higher than those in the plasma.^40^ The formulation of PSC is mixed with hydroxy-β-cyclodextrin, which improves its antifungal activity and pharmacokinetics by enhancing its solubility and oral bioavailability^41^ as well as ITC.^42^ These results suggest that PSC with improved tissue migration may be a suitable therapeutic option for invasive *E. dermatitidis* infections.

Recently, amikacin liposomal inhalation suspension (ALIS)^43^ was developed for the treatment of refractory nontuberculous mycobacterial infectious pulmonary disease (NTM-PD) and demonstrated better efficacy in the CONVERT trial.^44^ Direct administration at the site of infection, such as inhalation, can increase drug concentrations at the tissue level, resulting in greater antifungal efficacy than that of systemic administration, and leading to a reduction in the side effects of the drug. Inhalable and spray-dried microparticles of TRB^45,46^ have been studied for the treatment of pulmonary fungal infections to enhance the beneficial effects of antifungal drugs as well as AmB.^47^ TRB, which does not require particularly prolonged exposure (Fig. 4), would be a suitable inhaled drug for the treatment of *E. dermatitidis* pneumonia, which has been reported in approximately 6 % of patients with bronchiectasis and cystic fibrosis.^5^ In addition, TRB may be a promising combination drug (synergistic or indifferent effects *in vitro*) with azoles, including PSC, for the treatment of invasive *E. dermatitidis* infections.

In conclusion, TRB could become an even more useful and attractive antifungal drug if new routes of administration, such as inhalation or novel drug delivery systems, are developed to enhance TRB tissue migration.

## Acknowledgments

We thank the bacteriological examination staff of the Department of Laboratory Medicine at Kumamoto University Hospital.

## Funding

This work was supported by the Japan Society for the Promotion of Science, KAKENHI grant number JP21K16324 (T.N.), and a grant from Kobayashi Foundation, Kobayashi pharmacy related grant (H.N.).

## Transparency declarations

None to declare

## Author contributions

T.N. and T.Y. designed the research and, T.N., T.Y., and M.O. performed all the experiments. T.Y., D.M., and H.N. discussed the data and supported the preparation of the research. Y.J. and Y.T. supervised the personnel and the study. T.N. and H.N. obtained the necessary funding. T.N. and T.Y. wrote the manuscript, and T.Y., D.M., Y.J., and Y.T. advised or edited the manuscript. All authors read, commented on, and approved the final manuscript.

## Supplementary data

Fig.S1 to S5 and Table S1 to S3 are available as Supplementary data at JAC Online.

## Notes

### Competing Interest Statement

The authors have declared no competing interest.

