## Supplemental file for "Antifungal potency of terbinafine as a therapeutic agent against *Exophiala dermatitidis in vitro*"

### Fig. S1 *E. dermatitidis* information

**A**

| ITS | | D1/D2 | |
|---|---|---|---|
| Fungus | Identities (%) | Fungus | Identities (%) |
| *Exophiala dermatitidis* | 100.00 | *Exophiala nigra* | 99.34 |
| *Capronia munkii* | 91.07 | *Exophiala oligosperma* | 99.12 |
| *Capronia mansonii* | 91.07 | *Exophiala exophialae* | 98.90 |
| *Exophiala nidicola* | 90.08 | *Exophiala jeanselmei* | 98.68 |
| *Exophiala heteromorpha* | 89.81 | *Exophiala bergeri* | 98.87 |
| | | ⋮ | |
| | | *Exophiala dermatitidis* | 96.90 |

**B**

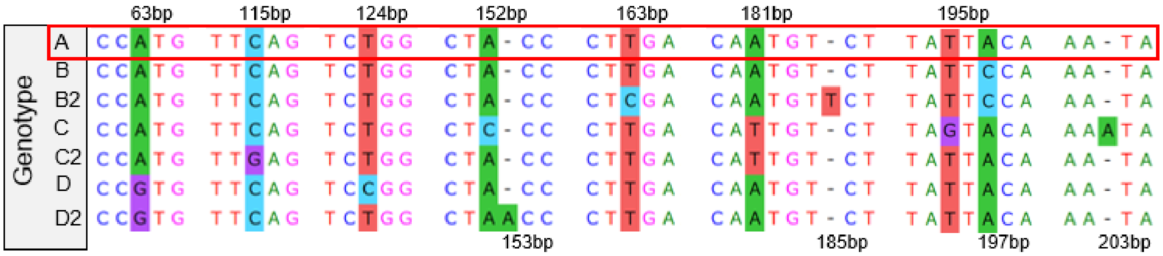


**C**

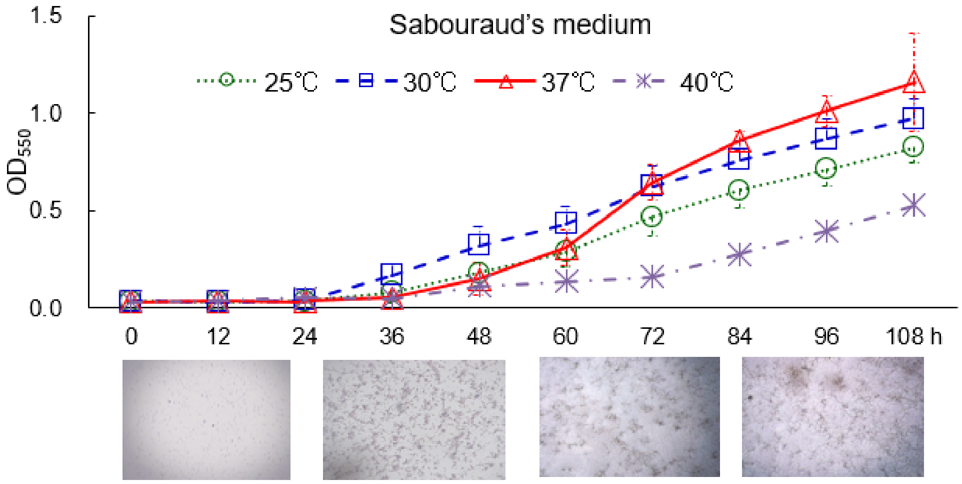


**A**) Genetic identification of *E. dermatitidis* from the ribosomal DNA (ITS and D1/D2). **B**) The DNA sequences of *E. dermatitidis* Genotype A, B, B2, C, C2, D, and D2. **C**) Growth rate and microscopic images of *E. dermatitidis* in Sabouraud's medium for 108 hours at 25, 30, 37, and 40°C.

### Fig. S2 SQLE 3D structures and drug chemical structures.

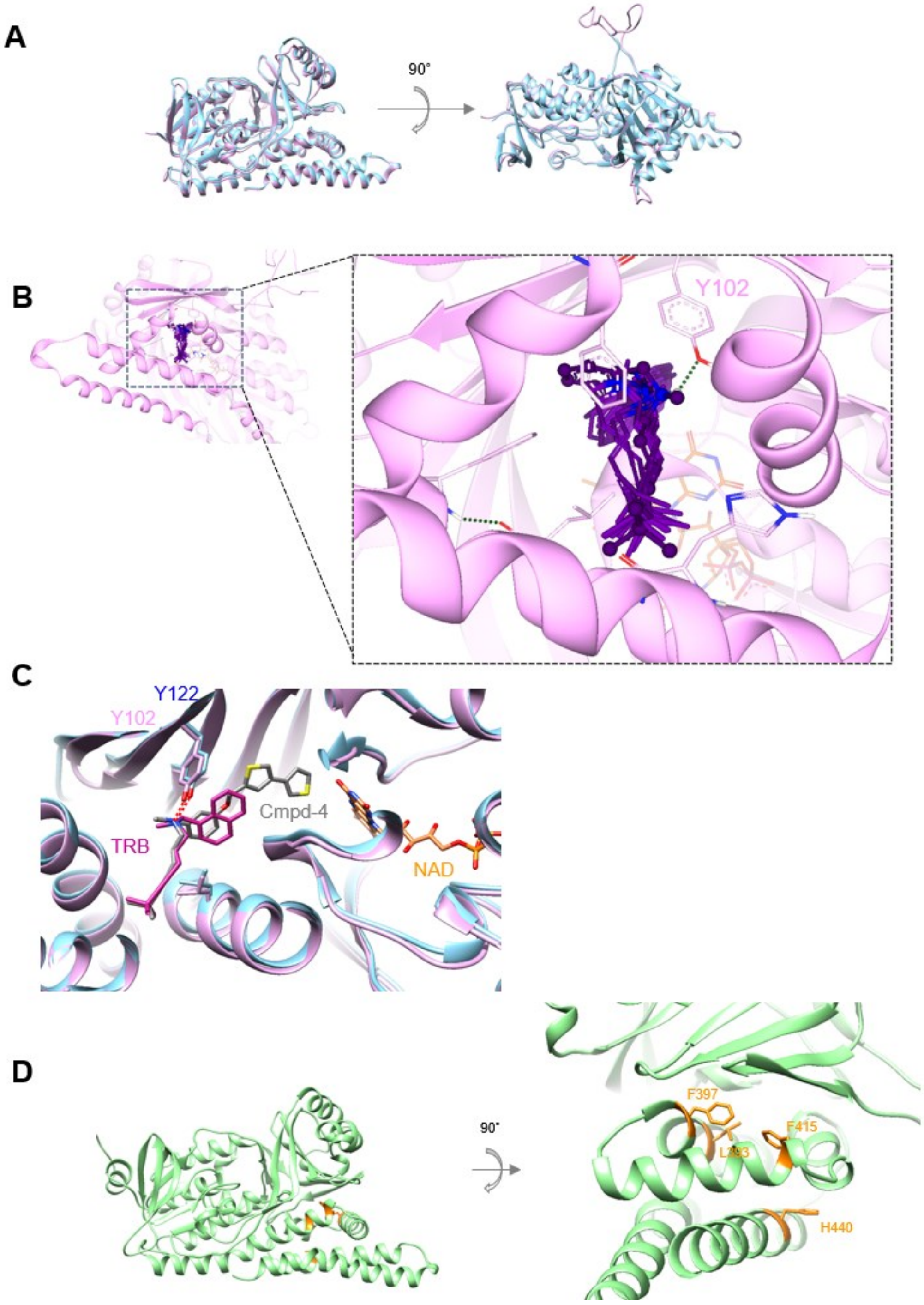

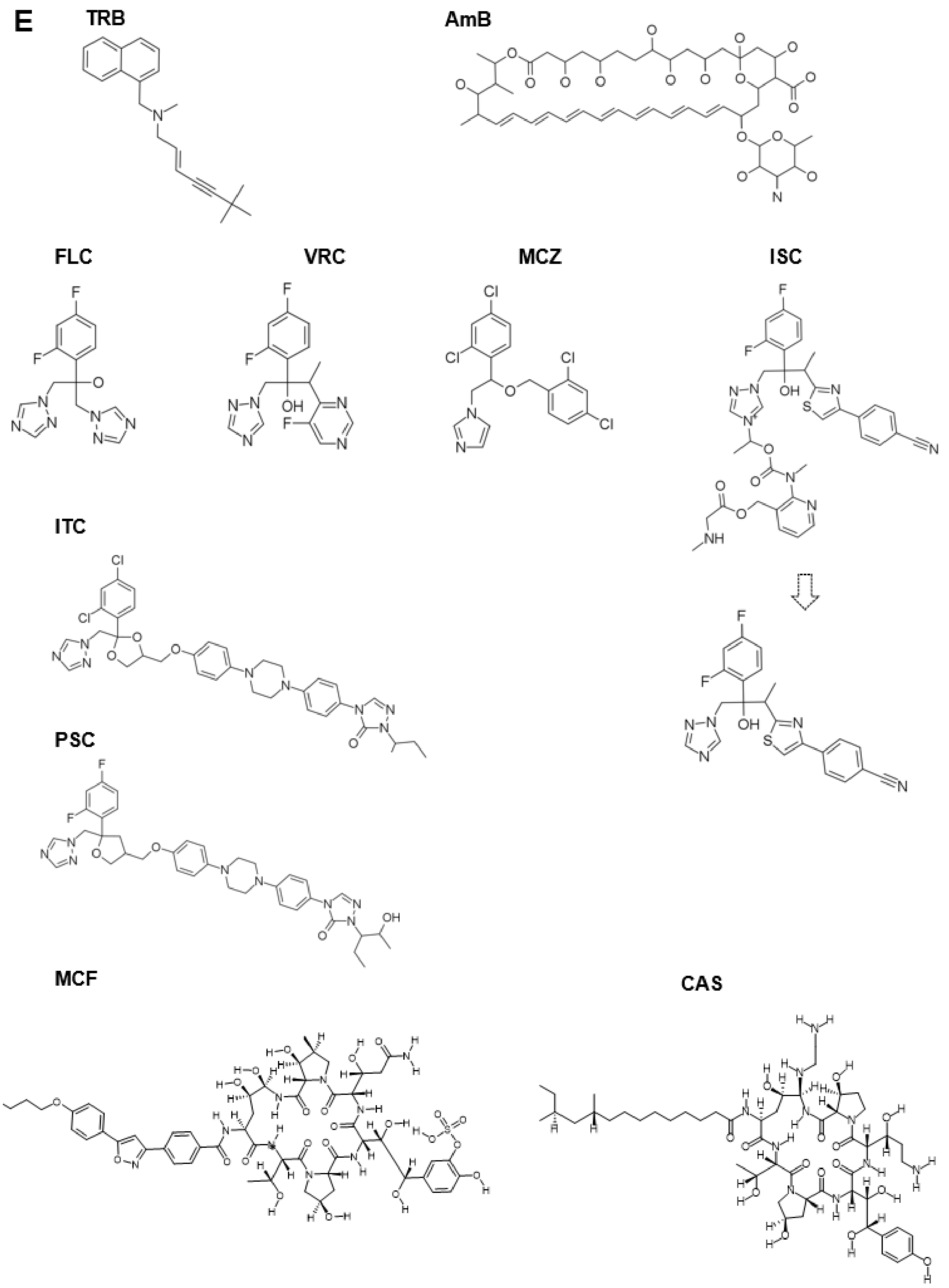


**A**) SQLE$^{Hum}$ and SQLE$^{ED}$ 3D structures are superimposed. **B**) The top 20 TRB binding to

SQLE$^{ED}$ modelling are shown. The binding pocket was defined as an area consisting of hydrophobic amino acid residues including Y102, L393, F397, F415, and H440. **C**) The 3D structure of the best TRB binding model to SQLE$^{ED}$ compared to cmpd-4 to SQLE$^{Hum}$ (6C6N). The H-bond interactions with Y102 and Y122 of SQLE$^{ED}$ and SQLE$^{Hum}$ respectively are shown as red dotted lines. **D**) The SQLE$^{TR}$ 3D structure modelling of the amino acid sequence of ATCC strain MYA-4607 is produced as a template for the crystal structure of SQLE$^{Hum}$ (PDB 6C6N) using the SWISS model. **E**) Chemical structures of all drugs used in this study using ChemDraw or MarvinSketch.

### Fig. S3 SEM observation of *E. dermatitidis* with or without A549 cells.

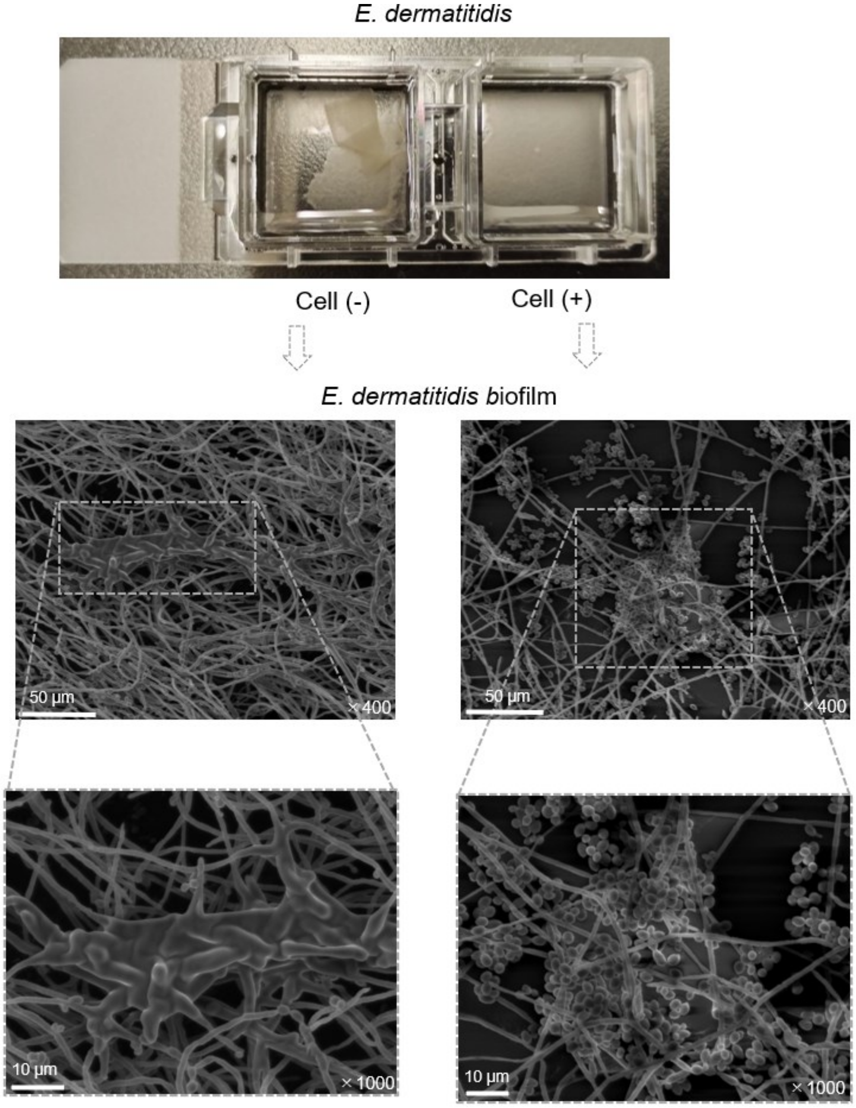

This image shows *E. dermatitidis* incubated for 48 hours at 35°C without A549 cells on the left side and with A549 cells on the right side in a coverslip set on a chamber slide II filled with DMEM containing 1% FBS, Cam, and K. These *E. dermatitidis* were observed to confirm the morphology including the biofilm by SEM. The scale bar and magnification at each image are shown.

### Fig. S4 Anti-biofilm and antifungal activity of TRB, PSC, AmB, and VRC against *E. dermatitidis*

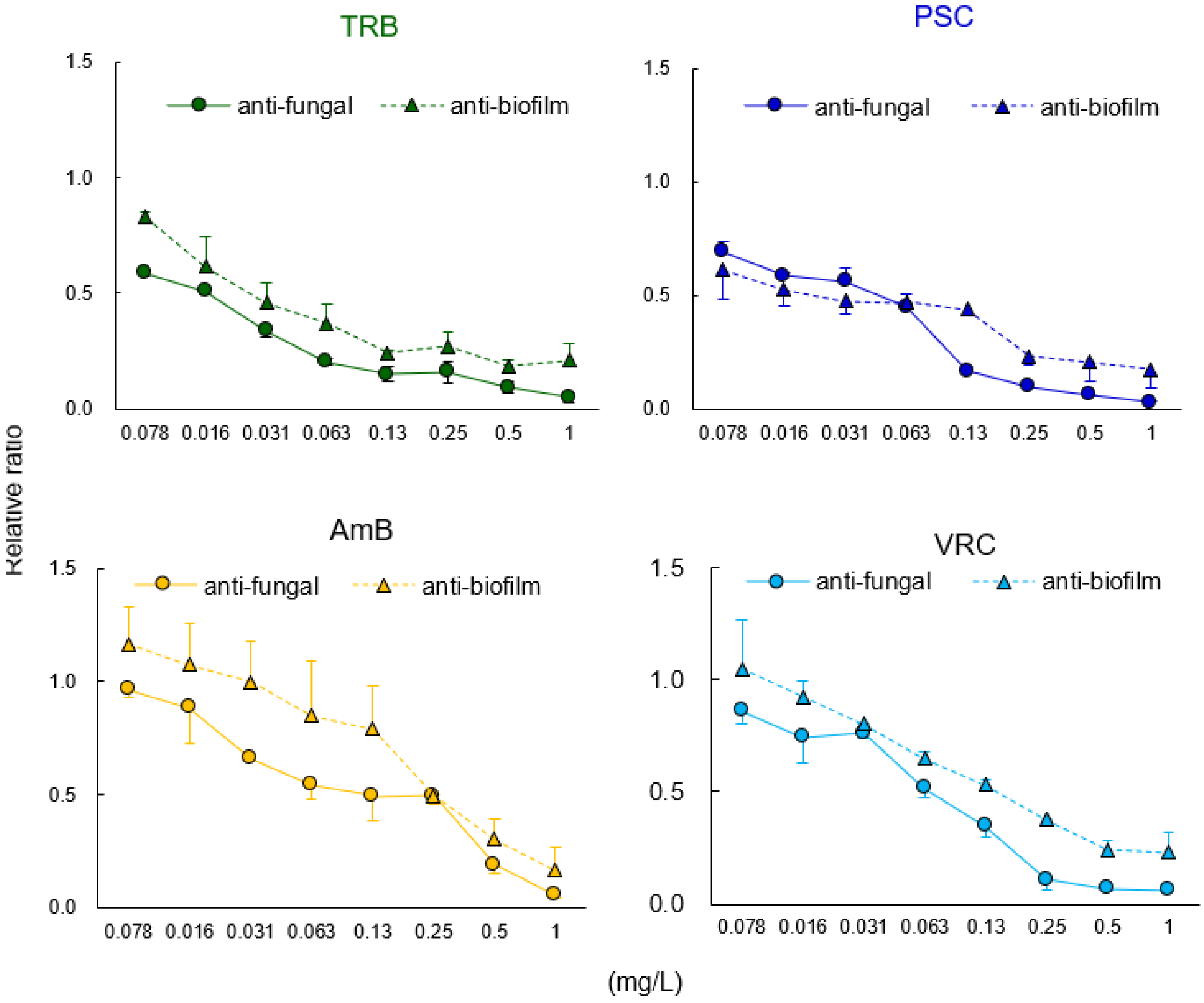


Anti-fungal activity of TRB, PSC, AmB, and VRC was determined by measuring *E. dermatitidis* growth at $OD_{530}$ before the CV assay was performed. All assays were performed independently in triplicate and the means (± S.D.) of representative data are shown.

**Fig. S5 Combination effect of TRB, ISC, ITC, PSC, and CAS on *E. dermatitidis***

## A

TRB + ISC

| Drugs | | TRB | | | | | | | | |
|---|---|---|---|---|---|---|---|---|---|---|
| | Conc. | 0.5 | 0.25 | 0.13 | 0.063 | 0.031 | 0.016 | 0.0078 | 0.0039 | 0 |
| ISC | 1 | 0.02 | 0.02 | 0.06 | 0.07 | 0.05 | 0.05 | 0.08 | 0.06 | 0.03 |
| | 0.5 | 0.01 | 0.01 | 0.05** | 0.18 | 0.26 | 0.45 | 0.40 | 0.38 | 0.46 |
| | 0.25 | 0.03 | 0.03 | 0.15* | 0.49 | 0.71 | 0.54 | 0.51 | 0.52 | 0.53 |
| | 0.13 | 0.01 | 0.01 | 0.14 | 0.17 | 0.57 | 0.82 | 0.93 | 0.83 | 0.84 |
| | 0.063 | 0.02 | 0.02 | 0.29 | 0.49*** | 0.76 | 0.85 | 0.76 | 1.00 | 0.51 |
| | 0 | 0.05 | 0.05 | 0.27 | 0.57 | 0.51 | 0.88 | 0.85 | 1.02 | 1.07 |

| | ISC | TRB | FICI |
|---|---|---|---|
| MIC* | 0.25 | 0.50 | 0.75 |
| MIC$_{90}$ WST-1** | 0.50 | 0.50 | 1.00 |
| MIC$_{50}$ WST-1*** | 0.13 | 0.50 | 0.63 |

0 - 0.1 | 0.1 - 0.5 | 0.5 -

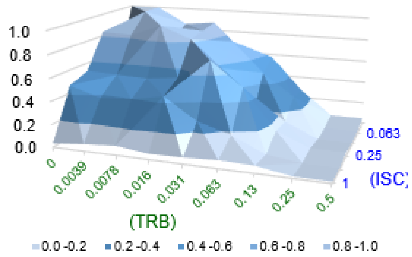


## B

TRB + VRC

| Drugs | | TRB | | | | | | | | |
|---|---|---|---|---|---|---|---|---|---|---|
| | Conc. | 0.5 | 0.25 | 0.13 | 0.063 | 0.031 | 0.016 | 0.0078 | 0.0039 | 0 |
| VRC | 0.5 | 0.04 | 0.04 | 0.04 | 0.04 | 0.04 | 0.04 | 0.04 | 0.04 | 0.03 |
| | 0.25 | 0.04 | 0.05 | 0.05 | 0.06 | 0.06 | 0.04 | 0.05 | 0.04 | 0.04 |
| | 0.13 | 0.02 | 0.04 | 0.03 | 0.07*/** | 0.15 | 0.22 | 0.16 | 0.24 | 0.30 |
| | 0.063 | 0.03 | 0.03 | 0.03 | 0.13 | 0.25 | 0.46 | 0.46*** | 0.52 | 0.57 |
| | 0.031 | 0.01 | 0.02 | 0.09 | 0.36 | 0.41 | 0.64 | 0.68 | 0.63 | 0.66 |
| | 0 | 0.02 | 0.02 | 0.08 | 0.42 | 0.61 | 0.86 | 0.79 | 0.81 | 0.91 |

| | VRC | TRB | FICI |
|---|---|---|---|
| MIC* | 0.5 | 0.5 | 1.0 |
| MIC$_{90}$ WST-1** | 0.5 | 0.5 | 1.0 |
| MIC$_{50}$ WST-1*** | 0.5 | 0.13 | 0.63 |

0 - 0.1 | 0.1 - 0.5 | 0.5 -

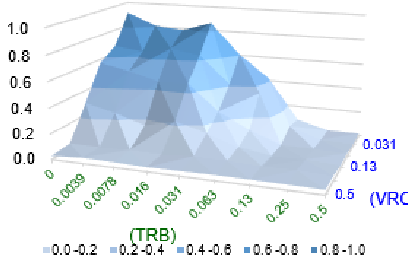


## C

TRB + ITC

| Drugs | | TRB | | | | | | | | |
|---|---|---|---|---|---|---|---|---|---|---|
| | Conc. | 0.5 | 0.25 | 0.13 | 0.063 | 0.031 | 0.016 | 0.0078 | 0.0039 | 0 |
| ITC | 0.5 | 0.03 | 0.02 | 0.02 | 0.03 | 0.02 | 0.01 | 0.01 | 0.04 | 0.01 |
| | 0.25 | 0.02 | 0.01 | 0.01 | 0.01 | 0.01 | 0.01 | 0.01 | 0.01 | 0.04 |
| | 0.13 | 0.01 | 0.01 | 0.02 | 0.06 | 0.12 | 0.09 | 0.13 | 0.12 | 0.15 |
| | 0.063 | 0.02 | 0.02 | 0.09 | 0.09*/** | 0.17 | 0.34 | 0.31 | 0.38 | 0.69 |
| | 0.031 | 0.02 | 0.07 | 0.20 | 0.20 | 0.49 | 0.44*** | 0.55 | 0.50 | 0.67 |
| | 0 | 0.03 | 0.03 | 0.12 | 0.27 | 0.59 | 0.86 | 0.67 | 0.79 | 0.95 |

| | ITC | TRB | FICI |
|---|---|---|---|
| MIC* | 0.25 | 0.25 | 0.5 |
| MIC$_{90}$ WST-1** | 0.25 | 0.25 | 0.5 |
| MIC$_{50}$ WST-1*** | 0.25 | 0.25 | 0.5 |

0 - 0.1 | 0.1 - 0.5 | 0.5 -

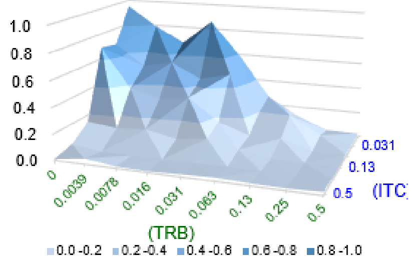


## D

CAS + PSC

| Drugs | | PSC | | | | | | | | |
|---|---|---|---|---|---|---|---|---|---|---|
| | Conc. | 0.5 | 0.25 | 0.13 | 0.063 | 0.031 | 0.016 | 0.0078 | 0.0039 | 0 |
| CAS | 32 | 0.01 | 0.01 | 0.01 | 0.01 | 0.01 | 0.02 | 0.03 | 0.02 | 0.04 |
| | 16 | 0.01 | 0.01 | 0.01 | 0.01 | 0.04 | 0.05 | 0.25 | 0.04 | 0.50 |
| | 8 | 0.01 | 0.01 | 0.01 | 0.07 | 0.14 | 0.49 | 0.44 | 0.46 | 0.67 |
| | 4 | 0.01 | 0.01 | 0.01 | 0.09* | 0.30 | 0.47*** | 0.50 | 0.78 | 0.73 |
| | 2 | 0.01 | 0.01 | 0.03 | 0.01** | 0.12 | 0.72 | 0.70 | 0.92 | 0.97 |
| | 0 | 0.00 | 0.01 | 0.09 | 0.52 | 0.68 | 0.69 | 0.71 | 0.78 | 0.87 |

| | CAS | PSC | FICI |
|---|---|---|---|
| MIC* | 0.13 | 0.25 | 0.38 |
| MIC$_{90}$ WST-1** | 0.06 | 0.5 | 0.56 |
| MIC$_{50}$ WST-1*** | 0.13 | 0.13 | 0.25 |

0 - 0.1 | 0.1 - 0.5 | 0.5 -

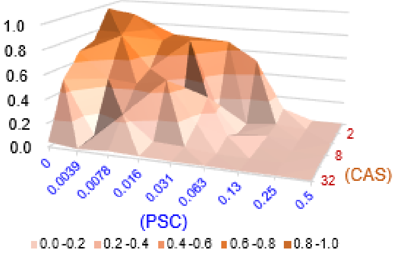

A combination of **A**) TRB and ISC, **B**) TRB and VRC, **C**) TRB and ITC, and **D**) CAS and PSC tables and 3D surface plots consist of relative ratios at each appropriate concentration as assessed by cell viability using WST-1 staining. FICIs were evaluated as synergy; FICI < 0.5, no interaction; 0.5 < FICI < 4, or antagonism; FICI > 4, which was calculated from MIC (*in red) data or $MIC_{50}$ (***) and $MIC_{90}$ (**) of WST-1 staining results. All assays were performed independently in triplicate and representative data are shown.

### Table S1 Amino acids sequences of SQLE among Humans, *T. rubrum (T.R.)*, *and E. dermatitidis (E.D.)*

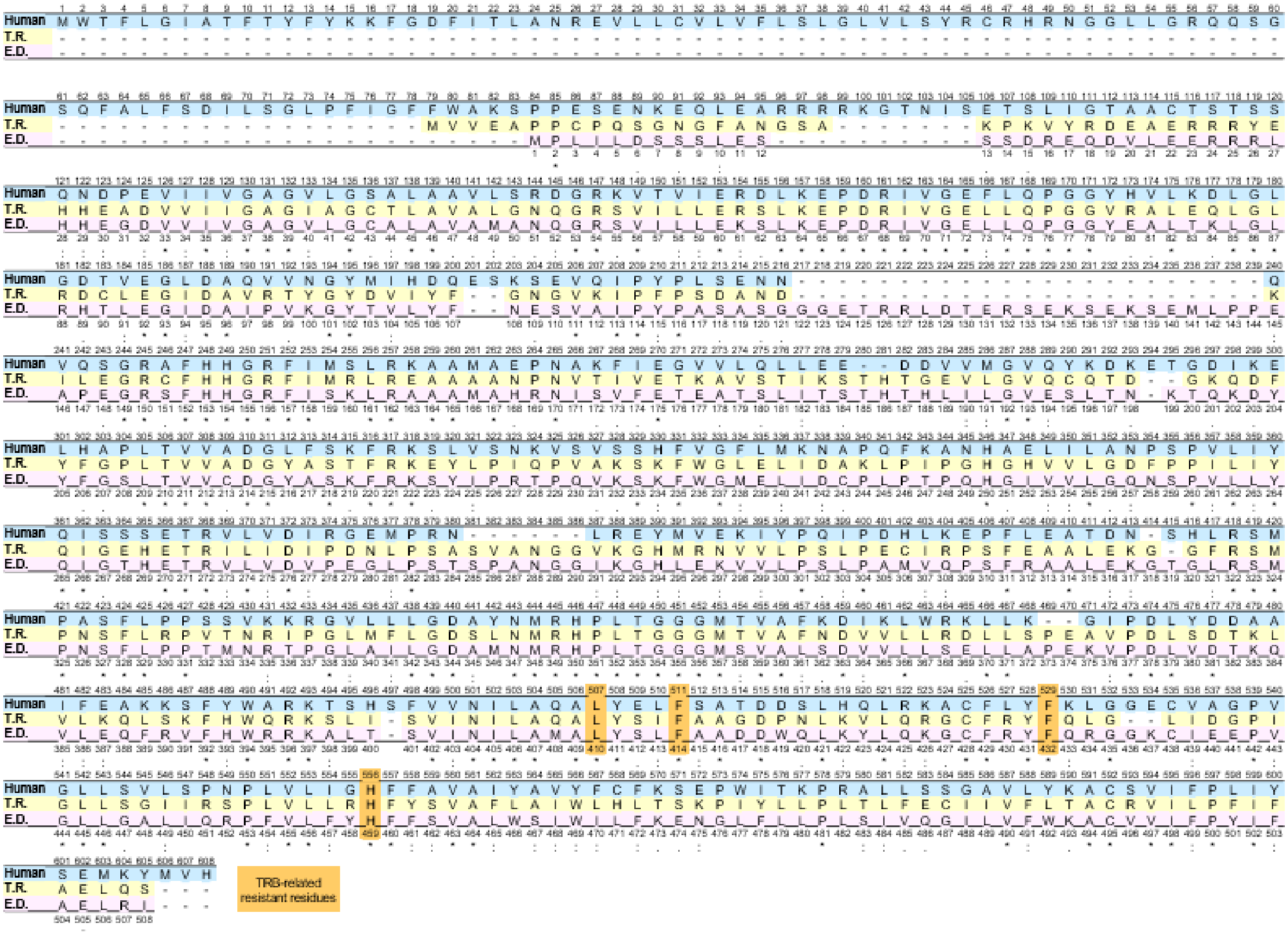


The amino acid sequences of SQLEs from Humans, *E. dermatitidis*, and *T. rubrum* were compared using Clustal omega (multiple sequence alignment tool; https://www.ebi.ac.uk/jdispatcher/msa/clustalo). TRB-related resistant amino acid mutations are shown in orange.

**Table S2 Susceptibility of *E. dermatitidis* 1, 2, and 3 to TRB, PSC, and AmB in this study**

| | | $OD_{530}$ | | $OD_{440}$ (WST-1) | | |
|---|---|---|---|---|---|---|
| **drugs** | **strains** | **$MIC_{50}$** | **$MIC_{90}$** | **$MIC_{50}$** | **$MIC_{90}$** | **MIC** |
| TRB | *E. dermatitidis. 1* | 0.13 | 0.25 | 0.125 | 0.25 | 0.25 |
| | *E. dermatitidis. 2* | 0.13 | 0.25 | 0.25 | 0.25 | 0.25 |
| | *E. dermatitidis. 3* | 0.13 | 0.25 | 0.5 | 0.5 | 0.25 |
| PCZ | E.D. 1 | 0.063 | 0.13 | 0.031 | 0.13 | 0.13 |
| | E.D. 2 | 0.063 | 0.13 | 0.13 | 0.25 | 0.13 |
| | E.D. 3 | 0.063 | 0.13 | 0.063 | 0.125 | 0.13 |
| AmB | E.D. 1 | 0.063 | 0.13 | 0.063 | 0.25 | 0.25 |
| | E.D. 2 | 0.13 | 0.5 | 0.25 | 0.5 | 0.5 |
| | E.D. 3 | 0.25 | 1 | 0.5 | 0.5 | 0.5 |

*E. dermatitidis 1* (E.D. 1) and *E. dermatitidis 2* (E.D. 2) were isolated from clinical patients in our hospital. *E. dermatitidis 3* (E.D. 3) (NBRC6421, ATCC28869) was purchased from NBRC. All *E. dermatitidis* are identified as genotype A. The susceptibility of all strains to TRB and PSC is the same, but that to AmB is different.

**Table S3 Anti-*E. dermatitidis* activity of the drugs at 25, 30, and 35℃**

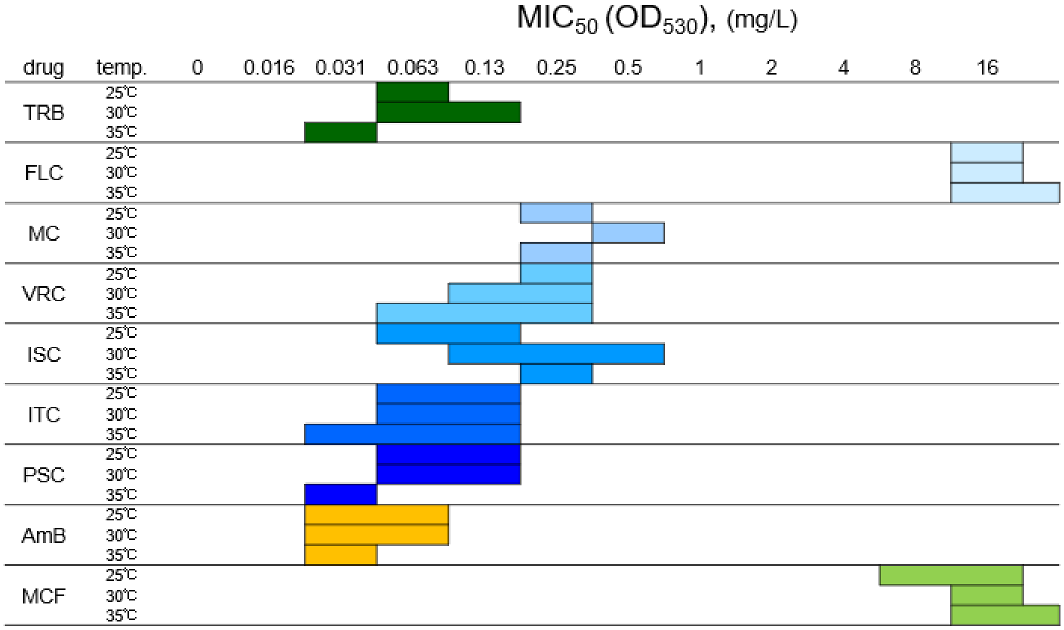


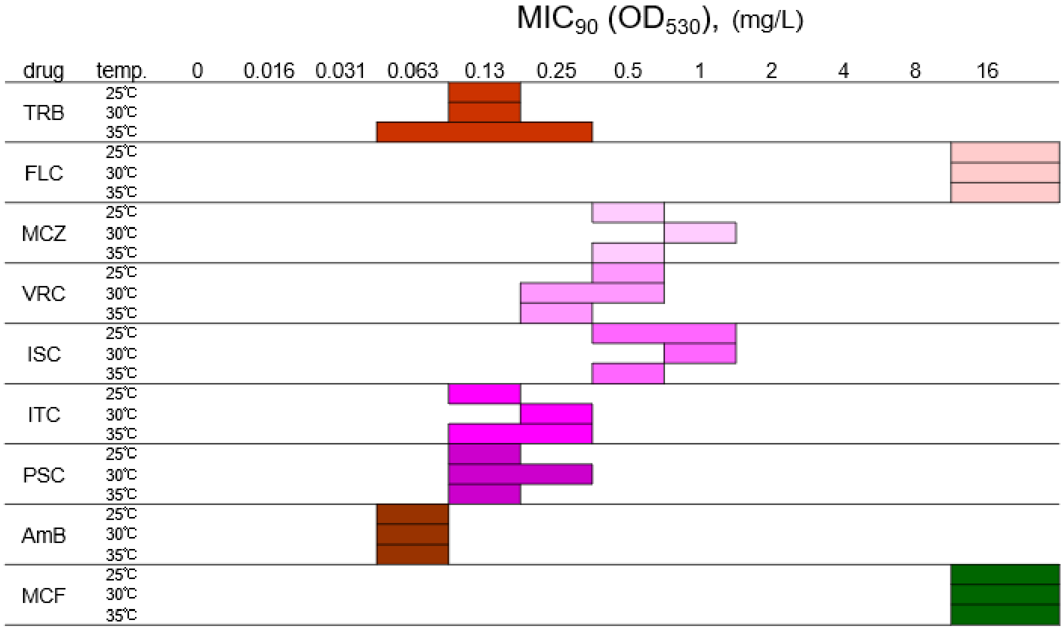


$MIC_{50}$ and $MIC_{90}$ of the drugs were determined by measuring *E. dermatitidis 1* growth at $OD_{530}$. All assays were performed independently in triplicate and all data are presented as ranges.
